# Exploring Chromosomal Structural Heterogeneity Across Multiple Cell Lines

**DOI:** 10.1101/2020.03.21.001917

**Authors:** Ryan R. Cheng, Vinicius Contessoto, Erez Lieberman Aiden, Peter G. Wolynes, Michele Di Pierro, José N. Onuchic

## Abstract

We study the structural ensembles of human chromosomes across different cell types. Using computer simulations, we generate cell-specific 3D chromosomal structures and compare them to recently published chromatin structures obtained through microscopy. We demonstrate using a combination of machine learning and polymer physics simulations that epigenetic information can be used to predict the structural ensembles of multiple human cell lines. The chromosomal structures obtained *in silico* are quantitatively consistent with those obtained through microscopy as well as DNA-DNA proximity ligation assays. Theory predicts that chromosome structures are fluid and can only be described by an ensemble, which is consistent with the observation that chromosomes exhibit no unique fold. Nevertheless, our analysis of both structures from simulation and microscopy reveals that short segments of chromatin make transitions between a closed conformation and an open dumbbell conformation. This conformational transition appears to be consistent with a two-state process with an effective free energy cost of about four times the effective information theoretic temperature. Finally, we study the conformational changes associated with the switching of genomic compartments observed in human cell lines. Genetically identical but epigenetically distinct cell types appear to rearrange their respective structural ensembles to expose segments of transcriptionally active chromatin, belonging to the A genomic compartment, towards the surface of the chromosome, while inactive segments, belonging to the B compartment, move to the interior. The formation of genomic compartments resembles hydrophobic collapse in protein folding, with the aggregation of denser and predominantly inactive chromatin driving the positioning of active chromatin toward the surface of individual chromosomal territories.

## Introduction

The 3D spatial organization of the chromosomes in the nucleus of eukaryotic cells appears to be cell type specific[1-7]. What determines this cell type specific organization and how that organization relates to patterns of gene expression remain crucial questions in structural genomics.

DNA-DNA ligation experiments have revealed spatial compartmentalization, generally termed A/B compartmentalization[8], and CTCF-mediated loop domains. It was observed that the A compartment chromatin contains a larger amount of the expressed genes while the B compartment chromatin is less transcriptionally active. Similar A/B compartmentalization has been observed across human cell lines[1-3] as well as in other species [2, 4, 9-12], suggesting that compartmentalization is a conserved feature of genome organization across evolution. While single-cell structures can be interrogated using proximity ligation assays[13, 14], high resolution has so far only been achieved through ligation methods when the experiments are performed over a large population of cells, thus averaging over the respective individual 3D structures.

Recent microscopy approaches have begun to reveal the 3D structures of segments of chromatin longer than a megabase at a spatial resolution on the nanometer scale[15-18]. These approaches not only allow for the quantification of pairwise and higher order interactions between loci, but also allow some quantification of the structural variability in a population of cells. One consistent observation from the imaging approaches, as well as from single cell Hi-C [13, 14, 19], has been the high degree of structural variability seen within an apparently homogeneous population of synchronized cells of a single cell type. Despite this variability, well-defined cell type-specific DNA-DNA ligation maps for the ensemble emerge after population averaging the single cell results.

Polymer models [20-29] that describe the process of chromosome organization have been proposed. In particular, the Minimal Chromatin Model (MiChroM) has been shown to accurately predict the population averaged DNA-DNA ligation maps[27, 30-32]. Chromosomes are described as polymers subject to interactions which depend on the chromatin biochemical composition and on the genomic distance separating any two loci[27]. Genomic distance-dependent interactions recapitulate the effect of motors acting along the DNA polymer and result in lengthwise compaction of chromatin. Interactions depending on chromatin biochemical composition recapitulate transient binding among chromosomal loci and result in the emergence of compartmentalization through a process of phase separation, in which chromatin of the same biochemical type preferentially co-localizes. The propensity toward phase separation for chromosomes of human lymphoblastoid cells can be reliably predicted using epigenetic marking data[30], suggesting that the information contained within the 1D epigenetic marking patterns decorating the chromatin polymer is sufficient to predict the ensemble of 3D chromosome structures. A neural network called MEGABASE[30] was trained to quantify the statistical relationship between the experimental sub-compartment annotations and the histone methylation and acetylation markings tracks, as assayed using chromatin immunoprecipitation data. Once trained, MEGABASE can be used to predict the compartmentalization patterns of a chromosome using a set of epigenetic ChIP-Seq tracks as the sole input. Combining MEGABASE and MiChroM, we are able to simulate the structural dynamics of chromosomes.

We first use the MEGABASE+MiChroM computational pipeline[30] to predict the 3D ensemble of chromosomal structures for several well-studied cell types: HMEC, HUVEC, IMR90, K562, HeLa-S3, and H1-hESC. To test these simulated 3D ensembles, we then generate ensemble averaged simulated ligation maps that are compared directly to population averaged DNA-DNA ligation maps[1, 2]. For the cell lines IMR90 and K562, we also use energy landscape tools to analyze the structures obtained through diffraction-limited microscopy by Bintu et al[15] for short ∼2 Mb segments of chromatin and compare the experimental structural ensembles directly with the corresponding regions of the simulated chromosome 21 for IMR90 and K562. This comparison shows that not only the population averages but also the structural heterogeneity that is observed in human chromosomes in the interphase are consistent with our energy landscape model. Chromosomes do not adopt a single structure in the interphase, but rather, exhibit a high structural variability characteristic of a phase-separated liquid. We provide a detailed characterization of this structural heterogeneity for the experimentally imaged and simulated segments of chromatin using an order parameter commonly used to quantify structural similarity in protein folding theory. For a gene-rich chromatin segment, we uncover two dominant clusters of structures in both the experimental and simulated structural ensembles: closed structures and open dumbbell-like structures. The transition from a closed structure to an open dumbbell appears to be governed by a two-state process with an apparent free energy cost of about four times the effective information theoretic temperature. For a gene inactive segment, structural analysis reveals highly disordered structures that lack domain boundaries. Additionally, we further examine the structural differences between whole chromosomes belonging to different cell types. The simulations show that inactive segments of chromatin move to the interior of the chromosome, while gene active chromatin moves to the chromosome surface. This effect appears to be driven by the favorable effective interactions between loci belonging to the B compartment, which forms a stable interior core; a phenomenon reminiscent of the hydrophobic collapse much studied in protein folding.

## Results & Discussion

### A polymer model of chromatin based on epigenetic features captures chromosome organization across different cell types

We previously developed a computational pipeline that can predict the 3D ensemble of chromosome structures by using chromatin immunoprecipitation tracks for histone modifications as input [30]. This approach was successfully used to predict the 3D chromosome structures for human lymphoblastoid cells (GM12878) using the experimental ChIP-Seq tracks for 11 histone modifications[30], i.e., H2AFZ, H3K27ac, H3K27me3, H3K36me3, H3K4me1, H3K4me2, H3K4me3, H3k79me2, H3K9ac, H3K9me3, and H4K20me1. Predicted chromosome structures for human lymphoblastoid cells (GM12878) were found to be consistent with both DNA-DNA ligation and fluorescence in situ hybridization (FISH) experiments[1]. Here we generate predictions beyond GM12878 to other well studied cell lines for which we have found sufficient epigenetic marking data.

Using the MEGABASE neural network, which was previously trained using data from GM12878, and sourcing from the Encyclopedia of DNA Elements (ENCODE) database the ChIP-Seq tracks for the same 11 histone modifications previously used, sub-compartment annotations for all the autosomes of cell lines were generated that had never been used in the training phase of the neural network. These sequences of sub-compartment annotations, or chromatin types, then serve as input for molecular dynamics simulations using the Minimal Chromatin Model (MiChroM)[27]. Using this combined approach, the chromosomal structural ensembles for 6 additional cell lines were generated: human fetal lung cells (IMR-90), human umbilical vein endothelial cells (HUVEC), immortalized myelogenous leukemia cells (K562), human mammary epithelial cells (HMEC), human embryonic stem cells (H1-hESC), and HeLa-S3 cells.

For each cell type, averaging the simulated ensemble generates *in silico* DNA-DNA ligation maps, which are in excellent agreement with those determined experimentally. Figure 1 shows the comparison between simulated and experimental maps for IMR90 (Figure 1A), HUVEC (Figure 1B) and K562 (Figure 1C), demonstrating quantitative agreement. Corresponding comparisons of the compartmentalization patterns are also provided in Figure S1 for additional cell types HMEC, H1-hESC, and HeLa-S3, as well as for GM12878 in Ref:[30]. In particular, the Pearson’s *R* between the simulated and experimental maps of matching cell type as a function of genomic distance shows that the long-range patterns of compartmentalization are captured over tens of mega-bases. To establish a term of comparison we calculated the Pearson’s *R* between the experimental DNA-DNA ligation maps of mismatching cell types. While the experimental observations on different cell lines do correlate with each other, computational modeling delineates the difference between cell type and appears to best match the experimental map when the cell types of simulation and experiment are matched up. This last result demonstrates that the present theoretical model discriminates well between different cell lines.

**Figure 1.**
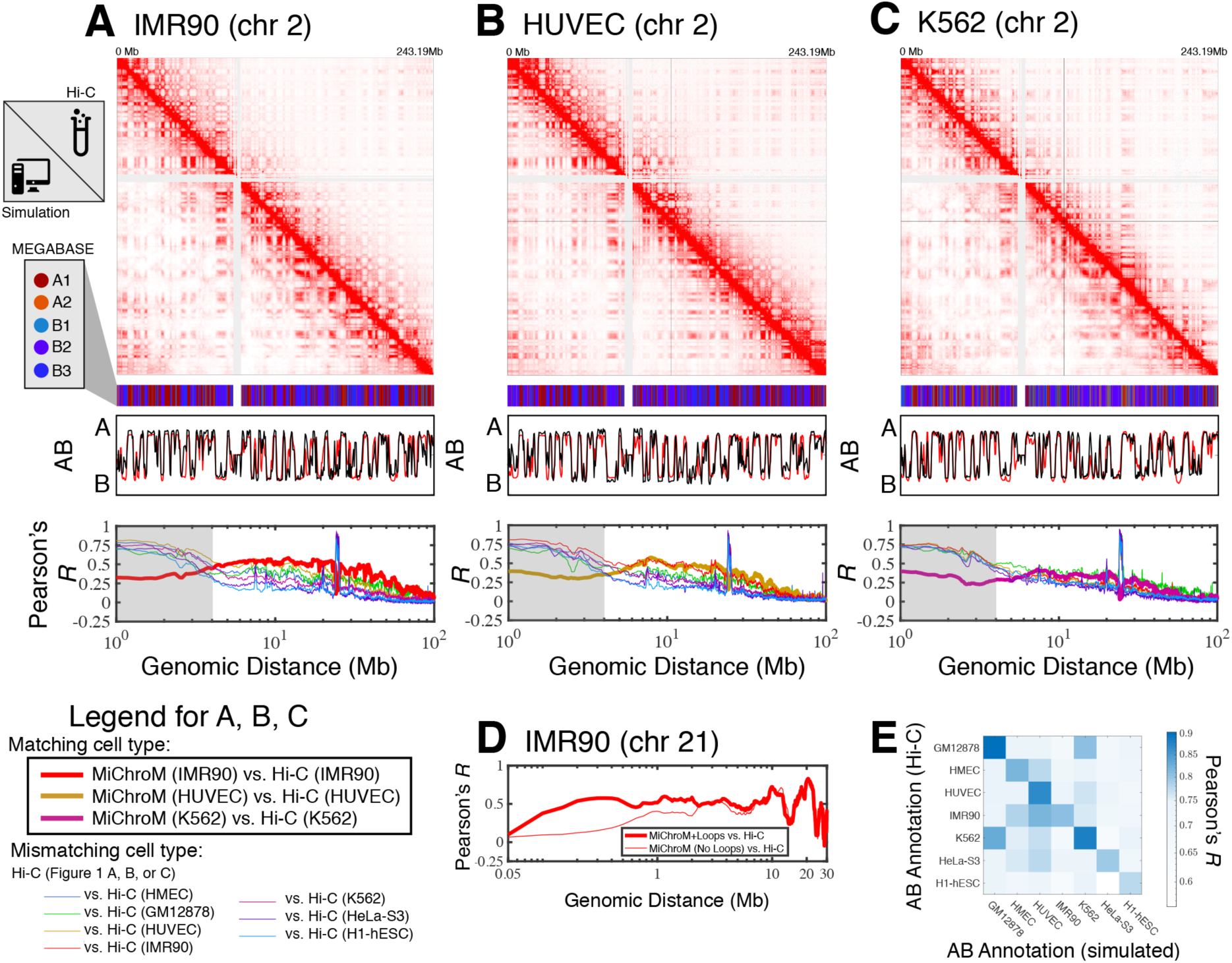
Prediction of chromosome structures for differentiated cell lines and for immortalized leukemia cells. The 3D ensemble of chromosome structures was predicted for the cell types (A) IMR90, (B) HUVEC, and (C) K562 using the ChIP-Seq histone modification tracks for the respective cell lines found on ENCODE—shown are the structural predictions for chromosome 2. As validation, the chromosome structures were compared with the DNA-DNA ligation experiments of Rao et al[1], where the simulated map is shown on the bottom left triangle and the experimental map is shown on the top right triangle. The datasets are visualized using Juicebox[33]. The MEGABASE chromatin type annotation is shown as a color vector under the contact probability map, followed by the A/B compartment annotation[1] for the simulated map (red) and the experimental map (black), respectively. The Pearson’s *R* between the simulated and experimental contact maps for fixed genomic distances are plotted for the cell types IMR90, HUVEC, and K562, respectively, in thick lines. The Pearson’s *R* between the experimental maps of mismatching cell types are also shown with thin lines—See Legend. The shaded region highlights that at relatively short genomic distances (<10 Mb), excluding CTCF-mediated loops from the simulation results in disagreement between the simulated and experimental maps. When loops are included in the simulations, the agreement between the simulation and experiment is recovered at the short genomic distances. (D) Pearson’s *R* as a function of genomic distance is plotted between the experimental map for chromosome 21 (IMR90) and MiChroM simulation with loops (thick red line) and without loops (thin red line). (E) A matrix of Pearson’s *R* between the AB annotation of the experimental ligation map and the simulated contact maps for different cell types, respectively. The high Pearson’s *R* signifies the consistency between the simulated maps and the experimental DNA-DNA ligation maps.

While we have focused so far on the spatial organization of entire chromosomes on the micrometer length scale, for a better comparison with the structures of chromosome 21 of IMR90 and K562 obtained from microscopy[15], we have also incorporated in the polymer physics simulation the loops mediated by the activity of the protein CTCF.

Figure 1D shows that the inclusion of CTCF-loops, which are easily be incorporated into the model, improves the quality of the results for the short range features of chromosome organization within 10Mb in genomic distance; at larger length scales the model appears to be completely insensitive of CTCF-mediated loops.

Figure 1E shows the Pearson’s *R* between the AB annotation vectors derived directly from the DNA-DNA ligation maps and those obtained from MiChroM simulations for different cell types. The diagonal of Figure 1E corresponds to the Pearson’s *R* between AB annotations derived from experiment and simulation of matching cell types. The simulated and experimental annotations for the same cell types agree well with each other. Figure S1E shows the Pearson’s *R* between AB annotations derived from experiment alone for the different cell types. Notably, the high degree of correlation between the myelogenous leukemia cell line K562 and human lymphoblastoids (GM12878) maps observed in Figure 1E is apparent from DNA-DNA ligation maps alone (Figure S1E). The agreement between the simulated and experimental A/B annotations is the highest quality (Pearson’s *r* ∼ 0.9) for the DNA-DNA ligation maps of GM12878, which is not surprising since the GM12878 has an order of magnitude more reads than any other map and consequently has the highest resolution.

Taken together, these results demonstrate that long range compartmentalization observed in the DNA-DNA ligation maps is well captured by the simulated structural ensembles for these well-studied cell lines using only information about the epigenetic marking patterns as input.

### Chromatin structural ensembles from DNA tracing reveal coexistence of open and closed structures

Recent developments in DNA-tracing have allowed the direct experimental determination of three-dimensional structures using diffraction-limited and super-resolution microscopy[15-18]. DNA tracing is a technique that labels consecutive stretches of DNA with optical probes, which can be used to spatially resolve the positions of those probes using microscopy. It has become increasingly clear that unlike the situation for folded globular proteins, which typically can be reasonably well approximated for many purposes by a single native structure corresponding to the average conformation, chromatin appears to be highly dynamical and cannot be characterized by any single conformation. The heterogeneity of the chromosomal structural ensembles was first suggested by the analysis of the free energy landscape of chromosomes[26, 27] and has been indirectly observed through single cell Hi-C experiments[13, 14, 19]. The heterogeneity has now been confirmed by direct imaging of individual chromosomal structures[15-17]. As a consequence of this conformational plasticity, statistical ensembles [26, 27, 30, 31, 34-39] must be used in order to describe chromosomal structures *in vivo*.

In order to improve our understanding of the genomic structural ensembles, we characterize the structural heterogeneity of chromatin that was imaged using microscopy. We focus on the traced structures of Bintu et al[15], who obtained hundreds of images structures for short ∼2Mb segments of chromatin belonging to chromosome 21. These regions are 29.37-31.32Mb (referred to here as Segment 1) of IMR90 and K562 cell types and 20.0-21.9Mb (referred to as Segment 2) of IMR90. Only structures where the positions of over 90% of the loci were resolved are used in the present analysis. There are then 692 usable structures for IMR90 Segment 1, 752 usable structures of IMR90 Segment 2, and 244 usable structures of K562 Segment 1.

As previously reported[15-17], the traced structures can be used to generate a population averaged contact maps, which turn out to be consistent with DNA-DNA ligation maps. Shown in Figure S2A-C are the averaged contact maps for the chromatin Segments 1 (IMR90 & K562) and Segment 2 (IMR90), respectively. Nevertheless, information is lost when converting from a 3D structural ensemble to a 2D contact map.

Focusing on the structural details that cannot be found in a contact map, we make a close examination of the types of structures observed in the tracing dataset using an order parameter commonly used in studying protein folding landscapes, *Q*, which quantifies the structural similarity between two structures α and β:

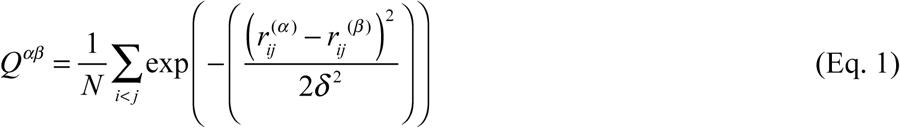

where 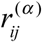 and 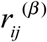 are the distances between chromatin loci *i* and *j* in structures α and β:, respectively, *N* is the number of pairs of loci included in the summation, and *δ* = 0.165*μm* is the resolution length scale for which deviations in the distances between structures α and β are treated as being similar. The *Q* between any two structures ranges from 0 (dissimilar) to 1 (identical) over the entire set of pairwise distances between loci. The order parameter *Q* is not solely based on contacts; a pair of chromatin loci can contribute to *Q* even if they are not spatially proximate if they are separated in both structures by a similar distance as set by δ. In this way *Q* measures structure more stringently than a simple contact map does.

Using 1− *Q* to define the distance between any two structures, hierarchical clustering of the traced structures for Segment 1 was applied to identify clusters having distinct structural features. These cluster sub-ensembles can be considered distinct conformational states. To see whether the segment 1 structures for IMR90 and K562 exhibit a high degree of structural similarity, we combined their datasets before clustering.

When applied to the 936 combined experimental structures for Segment 1, the clustering algorithm yields three distinct clusters. These correspond to a closed dumbbell (Cluster 1), an open dumbbell (Cluster 2), and a highly dense chromatin state (Cluster 3) shown in Figure 2. The closed dumbbell, where the head and tail globular domains are in contact with one another, is the dominant state observed for Segment 1 in both IMR90 and K562, accounting for 97.4% of the imaged structures (*N*_*closed*_ = 912). Cluster 1 can further be sub-divided into subgroups 1a, 1b, 1c, and 1d (Figure 2), which account for 75.5% of the structures in Cluster 1. The subgroups appear to capture various stages of the process of opening. The structures in subgroup 1a are fully collapsed, while structures in 1b, 1c, and 1d capture the progressive opening of the closed dumbbell. The distribution of the Radius of Gyration for structures belonging to sub-clusters 1a-1d is shown in Figure S3. The open dumbbell structures where the head and tail domains have dissociated from one another, account for approximately 1.8% of the imaged data (*N*_*open*_ = 17). Additionally, 7 dense, highly compact structures were identified from clustering. Representative structures from the three clustered structural groups are shown in Figure 2 and the corresponding population averaged contact maps are shown in Figure 2B and 2C for the closed and open structures, respectively.

**Figure 2.**
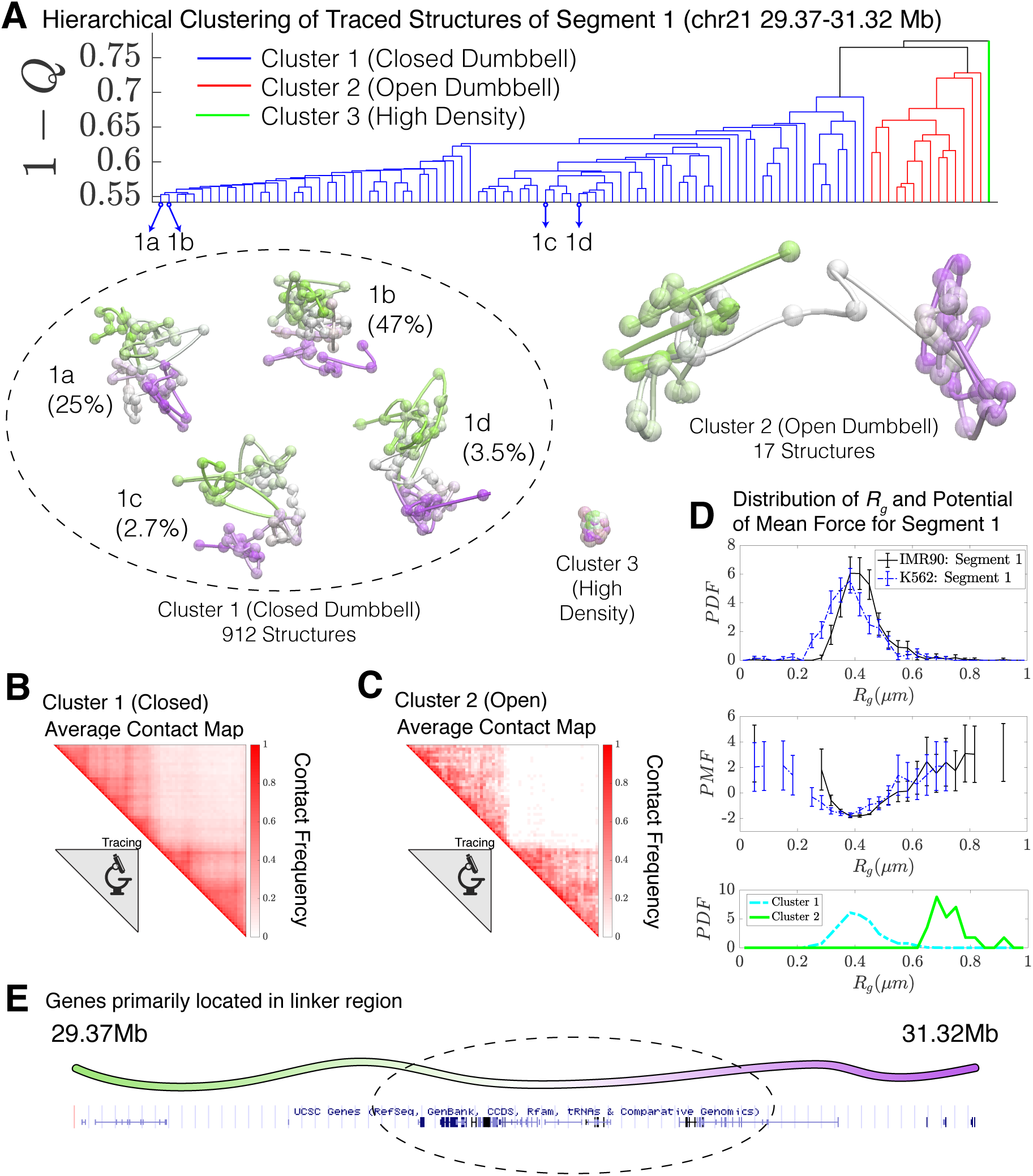
Hierarchical Clustering and the detailed structural analysis of traced Segment 1. (A) The dendrogram representation of the hierarchical clustering of Segment 1 (chr21 29.37-31.32Mb for IMR90 and K562 of [15]), where 1− *Q* is used as the distance between two structures. The clustering reveals three main clusters: closed dumbbell, open dumbbell, and highly dense structures. Further analysis of Cluster 1 reveals the presence of sub-clusters labeled 1a-1d that represent the gradual opening of the closed dumbbell. Representative traced structures are shown for each of the clusters and sub-clusters. The population averaged contact maps for the closed and open structure clusters are shown respectively in (B) and (C), where 165nm is used to define a contact between two 30kb loci. (D) The distribution of the Radius of Gyration (top), the corresponding potential of mean force (center), and the distributions of Radius of Gyration for Cluster 1 and Cluster 2 (bottom) are shown for the traced structures of Segment 1 of IMR90 and K562. The distribution exhibits a heavy tail to the right of the average value, indicating the existence of open, elongated structures. (E) The UCSC Genes track is plotted along the genomic positions of Segment 1 using the Genome Browser [42].

The high density chromatin, cluster 3, which was found when imaging both Segment 1 and Segment 2[15], is characterized by an extraordinarily high density of DNA ∼ 2 × 10^3^ *mg* / *ml*, as estimated for naked dsDNA. For comparison, the density of heterochromatin that is estimated using microscopy data is ∼ 200*mg* / *ml* [40]; for this reason, we believe that these chromatin conformations are likely artifacts of the experimental protocol. We therefore have excluded Cluster 3 from further analysis.

Assuming that the opening of the chromatin Segment 1 is in an effective thermodynamic equilibrium would imply a relative stability of log(*N*_*closed*_ / *N*_*open*_) = *E*_*open*_ − *E*_*closed*_ ∼ 4*k*_*B*_*T*, where ^*E*^_*open*_ ^− *E*^_*closed*_ is an apparent free energy difference between the closed and open states and *T* is an information theoretic temperature characterizing the ensemble[41]. Interestingly, the relative number of open and closed structures found in the simulations (discussed in the proceeding section) is in remarkable agreement with this experimental finding.

We then used the radius of gyration, *R*_*g*_, as an additional order parameter for the structural ensembles of segment 1 belonging to IMR90 and K562 (Figure 2D). A corresponding potential of mean force can be extracted from the distribution of *R*_*g*_ as *PMF* = −*k*_*B*_*T* log *P*(*R*_*g*_), which also shows the free energy difference of ∼ 4*k*_*B*_*T* between the closed (Cluster 1) and open (Cluster 2) structural sub-ensembles. The distributions of *R*_*g*_ are also shown for Clusters 1 and 2 in Figure 2D. The open conformations (Cluster 2) possibly belong to a free energy minima in the PMF located between between *R*_*g*_ ∼ 0.6 − 0.8*μm*, although additional statistics would be necessary to firmly establish the presence of this additional conformational mode. Interestingly, the vast majority of genes appear to be positioned along the linker region connecting the two globular domains (Figure 2E).

Unlike Segment 1, Segment 2 of IMR90 completely lacks loop domains and, consequently, the averaged contact maps for Segment 2 exhibit no additional features beyond the decay in contact probability as a function of genomic distance (Figure S2). Structural analysis reveals that, without the presence of loop domains, Segment 2 is highly disordered; while clustering reveals open and closed structures, the lack of loop domains and domain boundaries results in the loss of dumbbell-like structures (Figure S4). It should be noted that unlike Segment 1, Segment 2 has an absence of genes (Figure S7).

### The chromosomal structures obtained from physical modeling are consistent with those observed with microscopy

We compare the chromosome structures sampled in the simulations to the diffraction-limited microscopy structures of Bintu et al[15], finding that the conformational states observed using microscopy are also found in the simulated structural ensemble without any calibration or fine tuning of parameters. While MEGABASE+MiChroM, provides us with structures of entire chromosomes, we focus specifically on the same ∼2Mb chromatin segment within chromosome 21 for our direct comparison.

It is important to note that the simulated model, and the structural variability that it captures, was derived from the energy landscape learned from population-averaged DNA-DNA ligation data using the principle of maximum entropy[27]. MiChroM has been shown to be consistent with experimental ligation maps (Figure 1 and Refs:[27, 30, 32]), as well as the distribution of distances between Fluorescence in Situ Hybridization (FISH) probes[30] and several observations regarding chromatin dynamics[31].

Using the 1− *Q* as the distance between all simulated structures for Segment 1, we now performed hierarchical clustering of the simulated structures. The dendrogram of this clustering is shown in Figure 3A, which uncovers two main clusters in the structural ensemble: a closed dumbbell (Cluster 1) and an open dumbbell (Cluster 2). The closed and open structures are consistent with those observed in the Bintu et al[15] datasets. The representative structures of the closed and open conformations are shown in Figure 3, alongside the averaged contact maps for each of the clusters (Figure 3B-C), which are consistent with those determined experimentally (Shown in Figure 2B-C). The simulated Cluster 1 can again further be sub-divided into subgroups; 1α, 1β, 1γ, and 1δ represent the 4 most populated sub-groups (Figure 3), which comprise 66% of the simulated structures of Cluster 1. The subgroups appear to capture various stages of the process of opening. The structures in subgroup 1α are fully collapsed, while structures in 1β, 1γ, and 1δ capture the progressive opening of the closed dumbbell. The Radius of Gyration of sub-clusters 1α-1δ are shown in Figure S5.

**Figure 3.**
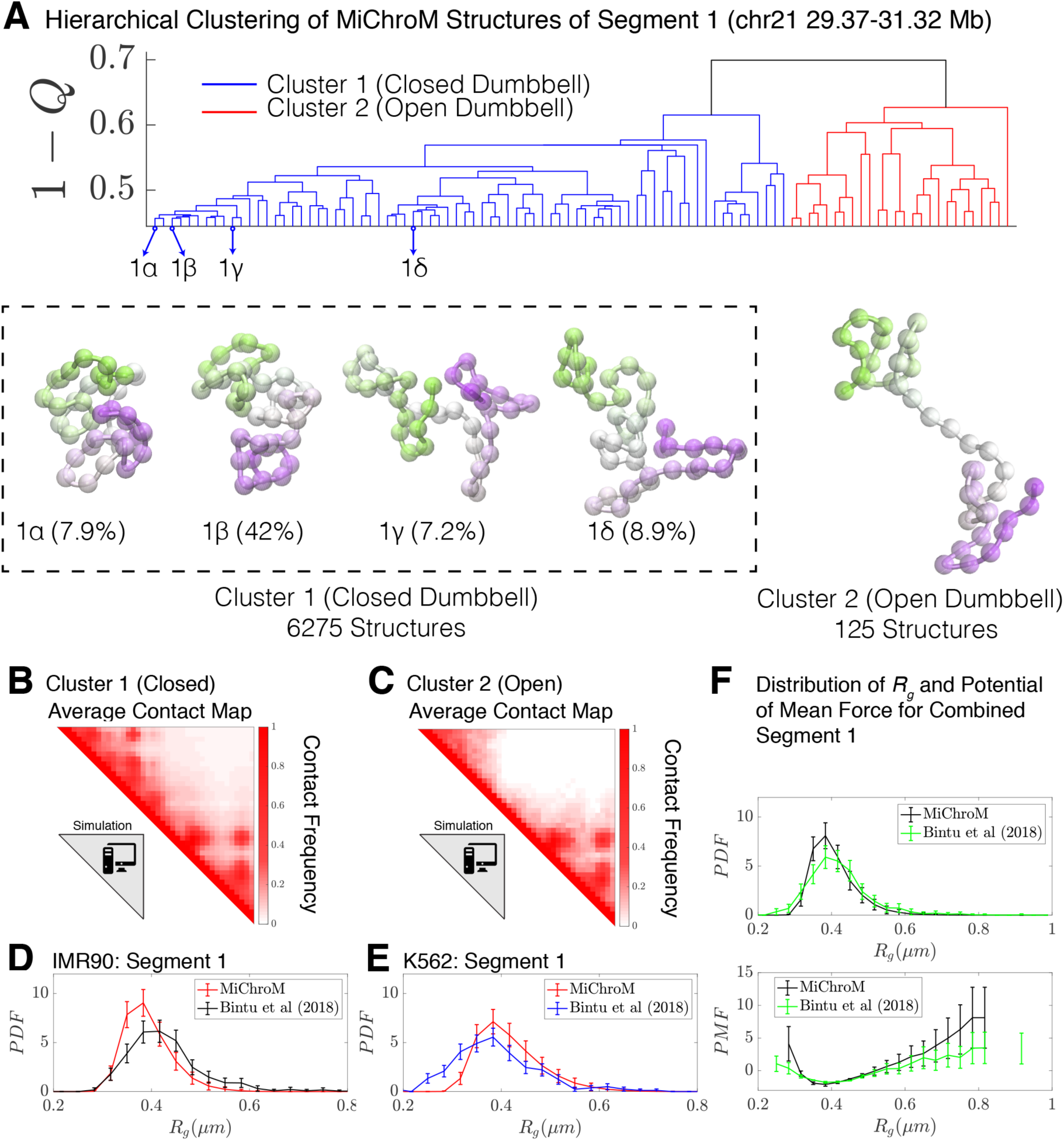
Hierarchical Clustering and the detailed structural analysis of simulated chromatin segment. (A) The dendrogram representation of the hierarchical clustering of simulated Segment 1 (chr21 29.37-31.32Mb for IMR90 and K562) where 1− *Q* is used as the distance between two structures. The clustering reveals two main clusters: closed dumbbell (6275 out of 6400 structures) and open dumbbell (125 out of 6400 structures). The closed dumbbell can be subdivided into sub-clusters labeled 1α-1*δ* that represent the opening transition of the closed dumbbell. Representative structures are shown for each of the clusters and sub-clusters. The population averaged contact maps for the clusters are shown respectively in (B) and (C), where 165nm is used to define a contact between two 50kb loci of the MiChroM model. The distribution of the Radius of Gyration is shown for Segment 1 IMR90 (D) and K562 (E) traced structures in comparison with the experimental structures. (F) Distribution of the Radius of Gyration and the corresponding potential of mean force is shown for both experiment and simulation for all of the structures of Segment 1.

No highly dense structures exist in the simulations. Such structures would collapse the entire chromatin segment to the volume of a single monomer, an occurrence that is prohibited by the energy function used to model the system. This is in harmony with our view that Cluster 3 seen in the experiments are artifacts of some sort.

For Segment 1, we performed our analysis on a set of 6400 structures, a representative subset of the simulated trajectories. Both closed (*N*_*closed*_ = 6275) and open structures *N*_*open*_ = 125) were identified by the clustering algorithm. Since MiChroM assumes an effective equilibrium thermodynamics representation of chromosome structures and dynamics, we can quickly calculate the relative stability between closed and open structures in the simulated ensemble as log(*N*_*closed*_ / *N*_*open*_) = *E*_*open*_ − *E*_*closed*_ ∼ 4*k*_*B*_*T*, where ^*E*^_*open*_ ^− *E*^_*closed*_ is the effective free energy difference between the closed and open states. This free energy difference is remarkably consistent with the value estimated using only the experimentally traced structures in the preceeding section.

Finally, we calculated the distribution of the radius of gyration, *R*_*g*_, for the experimetal traced structures of Bintu et al [15] and for the simulated MiChroM structures for Segment 1 belonging to IMR90 and K562 (shown in Figure 3D and Figure 3E respectively). Using a length scale calibrated previously[30] from a single FISH experiment of 0.165µm yields excellent quantitative agreement between the experimentally observed structures and those predicted *de novo* from simulation. It is particularly remarkable that any discrepancies between the experimental and simulated datasets can in fact be captured within 5% error of our original length estimate (Figure S6). Similarly, Figure 3F shows the direct comparison between the distribution of *R*_*g*_ for Segment 1 as well as the corresponding potential of mean force. We see then that MiChroM appears to reproduce the apparent free energy difference between open and closed structures found using the experimentally traced structures.

### Comparative analysis of the chromosomal structural ensembles of different cell lines: connecting the epigenetic markings of loci with their radial positioning within territories

The frequency of chromatin type annotations predicted by MEGABASE over different cell types is shown in Figure 4A as a stacked bar chart that represents the distribution of chromatin type annotations predicted for each locus of chromosome 2 over all of the cell types. It is evident that certain loci have similar epigenetic markings patterns in all the cell types that we examined, either by being generally transcriptionally active loci, thus likely belonging to the A compartment, or by being transcriptionally inactive B compartment loci. On the other hand, several segments of chromatin switch compartments between different cell types.

**Figure 4.**
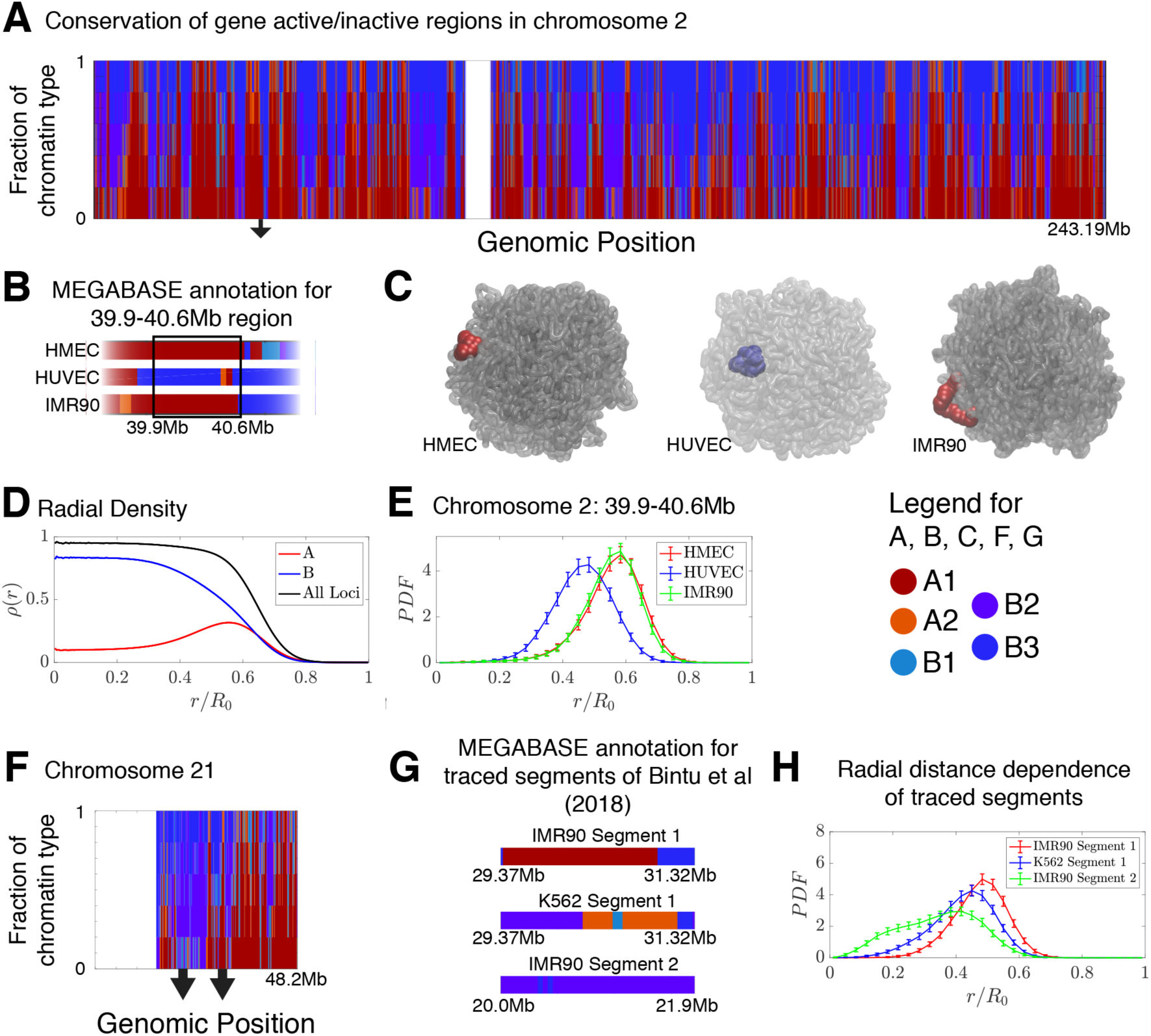
Conservation of compartmentalization across cell types and the radial dependence of marked chromatin. (A) A stacked bar chart is used to represent the distribution of chromatin type annotations predicted by MEGABASE as a function of the genomic position along chromosome 2 (hg19). The colors correspond the chromatin types given in the Figure Legend. For a given genomic position, the relative height of a particular color indicates the fraction of that particular chromatin type predicted at that locus. (B) The MEGABASE prediction of the chromatin type is shown for the chromatin segment 39.9-40.6 Mb of chromosome 2 for HMEC, HUVEC, and IMR90. A black arrow in (A) highlights the location of this segment. (C) The chromatin segment 39.9-40.6 Mb of chromosome 2 is shown in a representative structure for each of the cell types, where the color of the segment denotes its MEGABASE annotation. For HMEC and IMR90, the segment of chromatin tends towards the chromosome surface, whereas the segment tends towards the interior for HUVEC. (D) The radial density as a function of the normalized radial distance is plotted for A compartment loci, B compartment loci, and all loci. (E) The probability density functions of the radial distance are shown for the center of mass of the segment 39.9-40.6 Mb of chromosome 2 for HMEC, HUVEC, and IMR90, respectively. (F) A stacked bar chart is used to represent the distribution of chromatin type annotations predicted by MEGABASE as a function of the genomic position along chromosome 21 (hg19). The arrows indicate the locations of the traced segments of Bintu et al[15]: Segment 1 (29.37-31.32Mb) and Segment 2 (20.0-21.9Mb). (G) The MEGABASE annotation of the traced chromatin segments are given for IMR90 and K562. (H) The distribution of radial distances of the center of mass of each traced segment is shown.

Using the structural ensemble from the simulations based on the predicted compartments we then quantified the conformational differences between different cell types. On the chromosomal scale, structural differences emerge primarily from changes in the phase separation of epigenetically marked segments of chromatin. An example is illustrated in Figure 4B, which focuses on the region 39.9-40.6Mb of chromosome 2 for HMEC, HUVEC, and IMR90. The MEGABASE classification (Figure 4B) identifies the segment in HMEC and IMR90 as belonging to the A compartment, whereas the segment for HUVEC should belong to the B compartment. Representative 3D structures for this segment for each of the respective cell types are shown in Figure 4C.

A plot of the radial density of A compartment loci and B compartment loci is shown in Figure 4D. These radial densities are consistent with previously reported simulations[27]. Taking a look at the radial distance of the center of mass of the segment of chromatin in each of the cell types, one finds that the A compartment loci tend to localize towards the surface of the chromosome, while the B compartment loci of the HUVEC cell type tend to localize in the interior (Figure 4E). A similar positioning of transcriptionally active chromatin toward the periphery of chromosomal territories was also observed by Nagano et al[13] in mouse cells using Hi-C experiments.

We additionally use simulations to predict and examine the spatial positioning of the segments of chromatin examined by Bintu et al in the context of the entire chromosome 21. The experimental traced structures could not be used to ascertain the spatial positioning of those chromatin segments within chromosome 21 since only short segments were imaged rather than the entire chromosome. Figure 4F shows a stacked bar chart that represents the distribution of chromatin types predicted by MEGABASE for each genomic position of chromosome 21. Figure 4G shows the MEGABASE predictions for the traced segments, showing that IMR90 Segment 1 (29.37-31.32Mb) is composed of A-type chromatin while IMR90 Segment 2 (20.0-21.9Mb) primarily is composed of B compartment chromatin types. K562 Segment 1 (29.37-31.32Mb) appears to contain both A and B chromatin types. Figure 4H shows the radial distance distribution of the center of mass of these segments of chromatin, showing that IMR90 Segment 2 tends to be in the interior, IMR90 Segment 1 tends to lie near the chromosome surface, and K562 Segment 1 occupies an intermediate region.

The finding that there exists a radial ordering associated with the spatial compartmentalization is consistent with the fact that according to the MiChroM potential[27], contact interactions between B loci exhibit the most favorable energetic stabilization of all chromatin interactions. On the other hand, A to B or A to A type interactions are significantly weaker than the B to B interaction, but are comparably strong to each other (See Table S1). In other words, according to the MiChroM energetic parameters (which were originally learned from Hi-C maps), B loci drive the phase separation of the chromosomes. Much like a hydrophobic-polar model from protein folding, the B compartmentalization forms the stable core of the simulated chromosome and the weaker interactions between A compartment loci with A or B loci tends towards the surface, to minimize the free energy of the molecular assembly. Our theoretical model thus corroborates recent experiments[43, 44] that suggests heterochromatin phase separation is a major driving force behind genome organization, further highlighting the important role of phase separation in biological organization [27, 45-48].

## Discussion

DNA-tracing combined with diffraction-limited or super-resolution microscopy is beginning to shed light on the high degree of variability that is characteristic of chromosomal structures in the interphase[15-18]. These studies add to the growing body of evidence that a unique chromosomal fold simply does not exist in the interphase. Chromosome structures in the nucleus appear to be highly dynamical, owing to the many non-equilibrium processes in the cell, such as the activity of motor proteins.

The advances in genome imaging and the molecular simulation of chromosomes allows the development of order parameters able to quantify the structural similarities between different chromosome structures, and the degree of heterogeneity in the ensemble of structures. Our results demonstrate that the order parameter *Q*, commonly used in protein folding studies and structural biophysics, is a good order parameter for characterizing the structural ensemble of a segment of chromatin. Despite the high degree of conformational plasticity, it appears that for segments of chromatin as short as the ones imaged by Bintu et al [15] (∼2Mb in length), there do exist distinct clusters of chromatin structures that can be distinguished using the *Q* order parameter. The dominant structures found for chromatin Segment 1 (chr21 29.37-31.32Mb) examined using data from microscopy as well as from simulation can be described as being a closed dumbbell and an open dumbbell, where the ends of the dumbbell are the globular domains at the head and tail of the chromatin segment.

It is known that CTCF proteins bound along the genome acts as gene insulators, probably through their suppressing activity toward loop extrusion[49-51]. Interestingly, a survey of the positioning of genes along Segment 1 reveals that the vast majority of genes appear clustered in the linker region of the chromatin segment (Figure 2E; Figure S7A), sandwiched between the head and tail loop domains. On the other hand, there is an absence of genes located on Segment 2 (chr21 20.0-21.9Mb) (Figure S7B), which contains no loop domains. Classification of the experimentally imaged structures of Segment 2 lack the domain boundaries that segregate the head and the tail of the chromatin segment into globular domains, although still exhibiting open and closed conformations. These findings suggest a possible role in transcriptional regulation for the opening and closing of organized dumbbell structures. How open and closed structures would achieve such regulation of the transcriptional activity remains to be investigated. It is however clear that understanding the structure-function relationship in the genome is a crucial question that can only be answered using an accurate statistical characterization of the conformational ensembles.

Finally, our work refines the classical view of the spatial compartmentalization of chromatin. We find a striking dependence between radial positioning of chromatin and epigenetic marking patterns. Our theoretical model, MiChroM, predicts that transcriptionally active loci, typically belonging to the A compartment, move towards the surface of the chromosomal territory, while B compartment loci, typically inactive, move to the interior[27]. Since interactions among B-B loci result in the greatest energetic stabilization, aggregation of these loci seems to be driving force behind both the phase separation of epigenetically similar chromatin into compartments and the expulsion of the active chromatin toward the periphery of chromosomal territories. In other words, according to the present energy landscape model, when the epigenetic marking patterns of a locus are rewritten from A to B, the locus moves towards the interior of the chromosome, perhaps affecting the transcriptional activity of the associated genes.

### Notes

Unless explicitly stated otherwise, all genomic positions are reported using the positions of the hg19 assembly. All of the simulated chromosome structures discussed in this manuscript were deposited in the Nucleome Data Bank (NDB)[32] found at https://ndb.rice.edu.

## Acknowledgements

The authors would like to thank Ting Wu for helpful discussions. This work was supported by the Center for Theoretical Biological Physics sponsored by the National Science Foundation NSF Grant PHY-1427654. J.N.O. was also supported by the NSF-CHE-1614101 and by the Welch Foundation (Grant C-1792). J.N.O. is a CPRIT Scholar in Cancer Research sponsored by the Cancer Prevention and Research Institute of Texas. V.G.C. is a Robert A. Welch Postdoctoral Fellow and was also funded by FAPESP (São Paulo Research Foundation and Higher Education Personnel: Grant 2016/13998-8), and CAPES (Higher Education Personnel Improvement Coordination: Grant 2017/09662-7). Additional support to P.G.W. was provided by the D. R. Bullard-Welch Chair at Rice University (Grant C-0016). E. L. A. was also supported by an NIH New Innovator Award (1DP2OD008540-01), the NHGRI Center for Excellence for Genomic Sciences (HG006193), the Welch Foundation (Q-1866), an NVIDIA Research Center Award, an IBM University Challenge Award, a Google Research Award, a Cancer Prevention Research Institute of Texas Scholar Award (R1304), a McNair Medical Institute Scholar Award, an NIH 4D Nucleome Grant (U01HL130010), an NIH Encyclopedia of DNA Elements Mapping Center Award (UM1HG009375), and the President’s Early Career Award in Science and Engineering.

## Supporting Information

**Figure S1.**
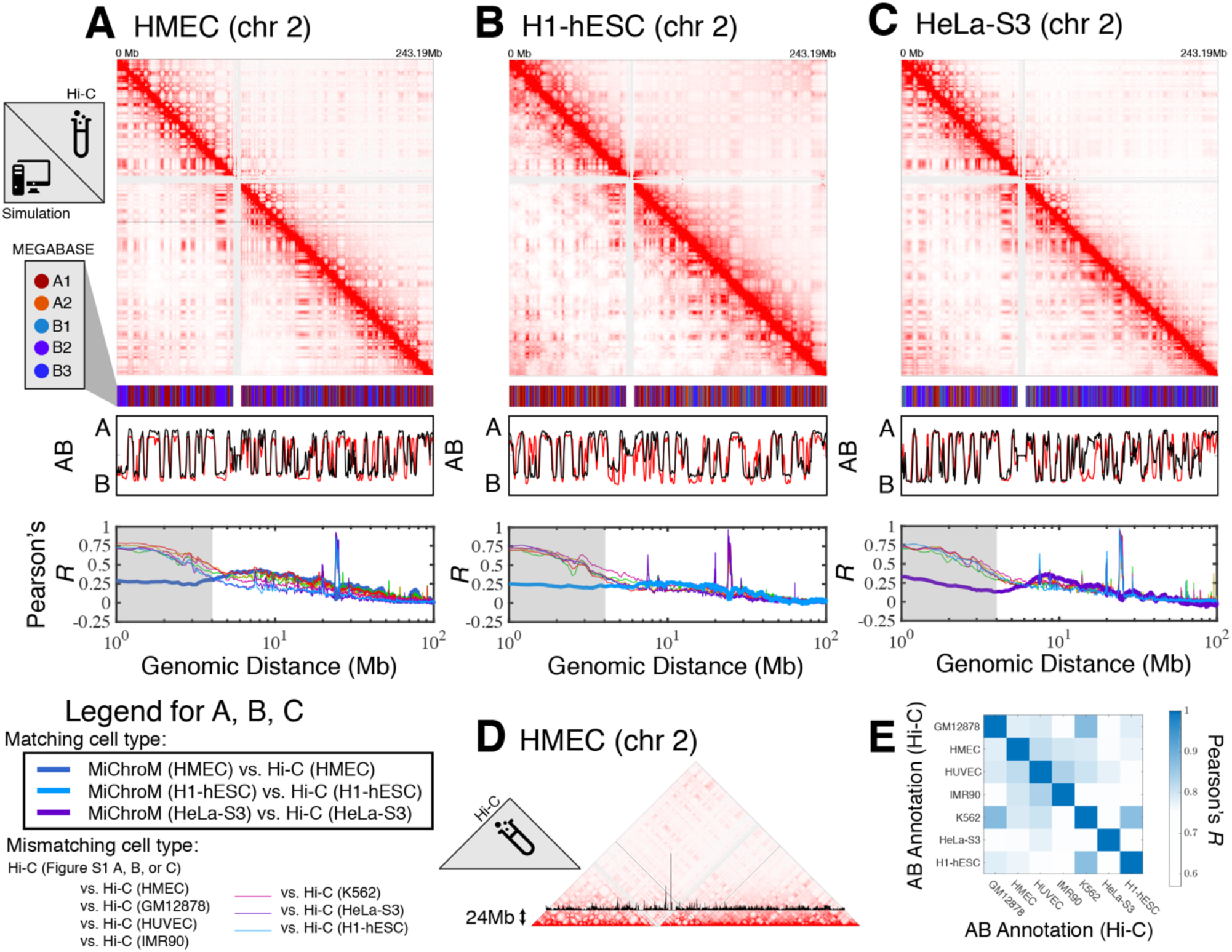
Prediction of chromosome structures for HMEC, H1-hESC, and HeLa-S3. The 3D ensemble of chromosome structures was predicted for the cell types (A) HMEC, (B) H1-hESC, and (C) HeLa-S3 using the ChIP-Seq histone modification tracks for the respective cell lines found on ENCODE—shown are the structural predictions for chromosome 2. As validation, the chromosome structures were compared with the DNA-DNA ligation experiments of Rao et al[1], where the simulated map is shown on the bottom left triangle and the experimental map is shown on the top right triangle. The datasets are visualized using Juicebox[2]. The MEGABASE chromatin type annotation is shown as a color vector under the contact probability map, followed by the A/B compartment annotation[1] for the simulated map (red) and the experimental map (black), respectively. The Pearson’s *R* between the simulated and experimental contact maps for fixed genomic distances are plotted for the cell types HMEC, H1-hESC, and HeLa-S3, respectively, in thick lines. The Pearson’s *R* between the experimental maps of mismatching cell types are also shown with thin lines—See Legend. (D) The experimental Hi-C map of HMEC (chr 2) [1] from (A) is shown while highlighting (in black) the relative magnitude of the contact probability along the strip corresponding to a fixed genomic distance of 24 Mb between pairs of loci. The large peak in the contact probability located proximal to the centromere and is present in all of the experimental Hi-C maps for IMR90, HUVEC, K562, HMEC, H1-hESC, HeLa-S3, and GM12878 and is responsible for the sharp peak in the Pearson’s *R* vs. genomic distance located at approximately 24 Mb in Figure 1A-C and Figure S1 A-C. The origin of this anomalous peak in the experimental map is unclear and it is not captured by the simulation. (E) A matrix of Pearson’s *R* between the AB annotation of the experimental ligation maps for different cell types. The diagonal elements of this matrix show the Pearson’s *R* of a particular AB annotation from Hi-C with itself and is equal to 1 by definition.

**Figure S2.**
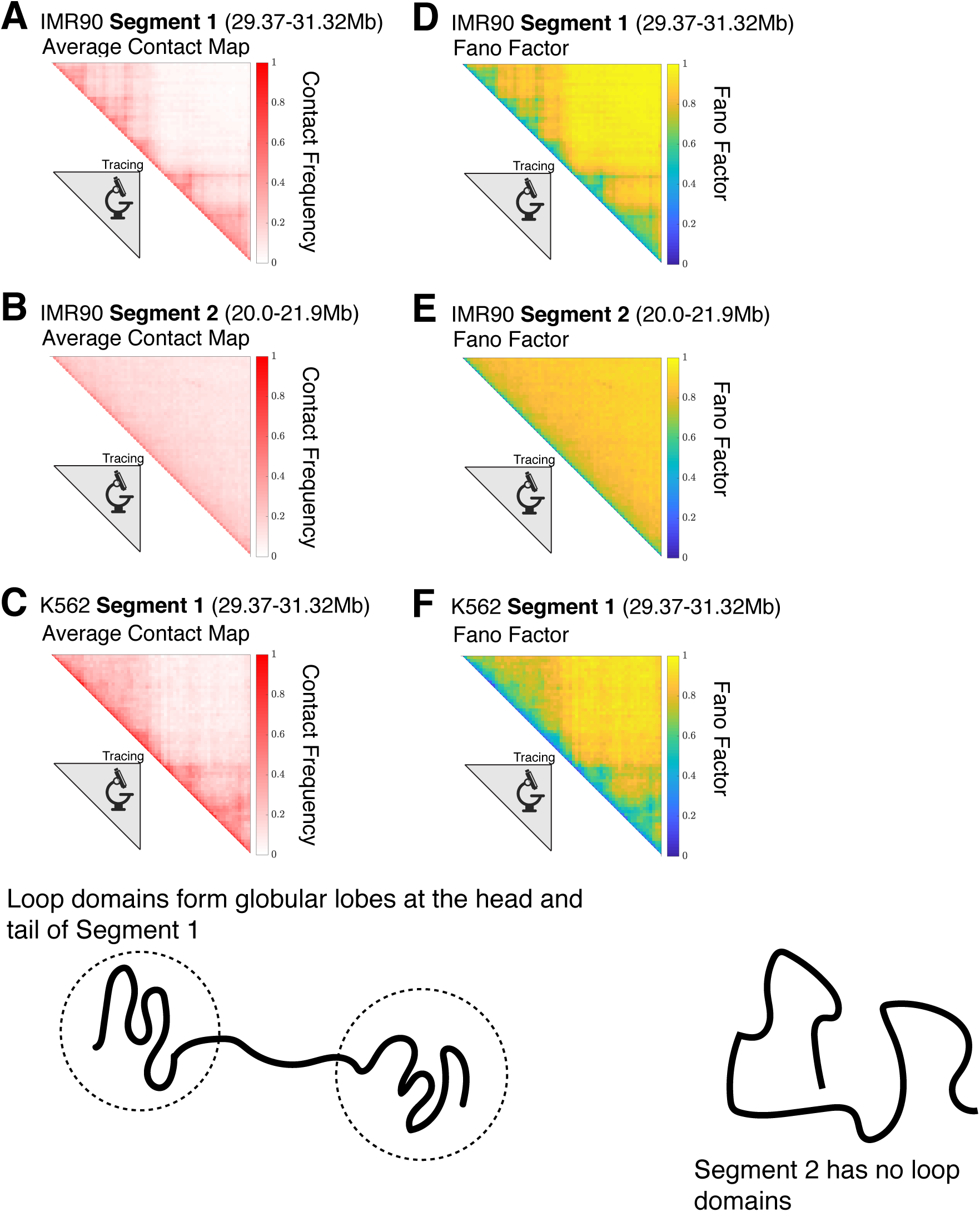
Contact maps for the experimentally traced segments of chromatin. The contact maps for the chromatin structures obtained from super-resolution imaging[3] for (A) IMR90 Segment 1 (chr 21 29.37-31.32 Mb), (B) IMR90 Segment 2 (chr 21 20.0-21.9 Mb), and (C) K562 Segment 1 (chr 21 29.37-31.32 Mb) are shown. For a given chromatin structure, two loci *i* and *j* are spatially proximal when the cartesian distance between them, *r*_*ij*_, is less than or equal to a cutoff distance *d—*we define a contact using *d* = 0.165*μm*. This allows us to define a label *c*_*ij*_ for a pair of loci that equals 1 when loci *i* and *j* are in contact and 0 when they are not: 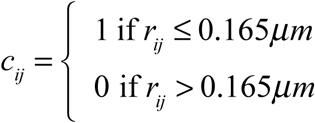. The variance of the contact frequency over the mean contact frequency (Fano Factor) *F* = (1− ⟨ *c*_*ij*_ ⟩) is plotted for each of the respective chromatin segments in (D), (E), and (F), where the angular brackets denote averaging over the structural ensemble. A Fano factor of *F* = 0 indicates zero variability, 0 < *F* < 1 indicates that the process is under-dispersed and characterized by a binomial distribution, and *F* = 1 is characteristic of a Poisson process. Segment 1 for IMR90 and K562 both have globular domains at the head and tail of the chromatin segment. On the other hand, Segment 2 has no loop domains and the only observable feature in its contact map is the decay of the contact probability as a function of genomic distance.

**Figure S3.**
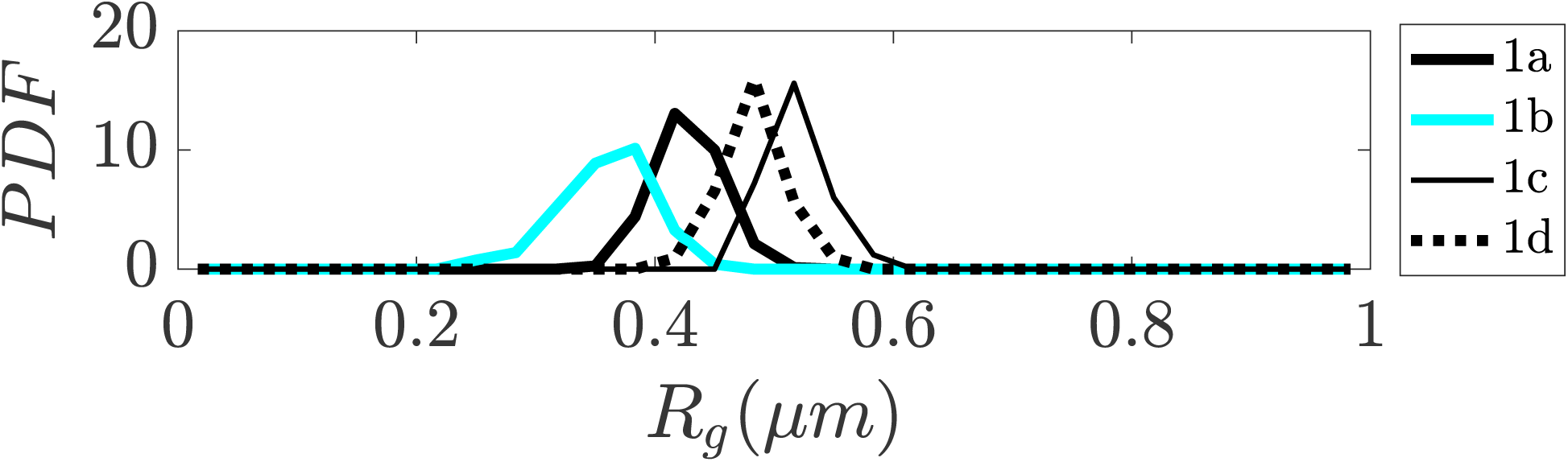
Distribution of Radius of Gyration for sub-clusters of closed dumbbell structures obtained experimentally using tracing. Sub-clusters of Cluster 1 in Figure 2 are denoted as 1a, 1b, 1c, and 1d. The sub-clusters of Cluster 1 characterize the gradual opening of the closed dumbbell structures.

**Figure S4.**
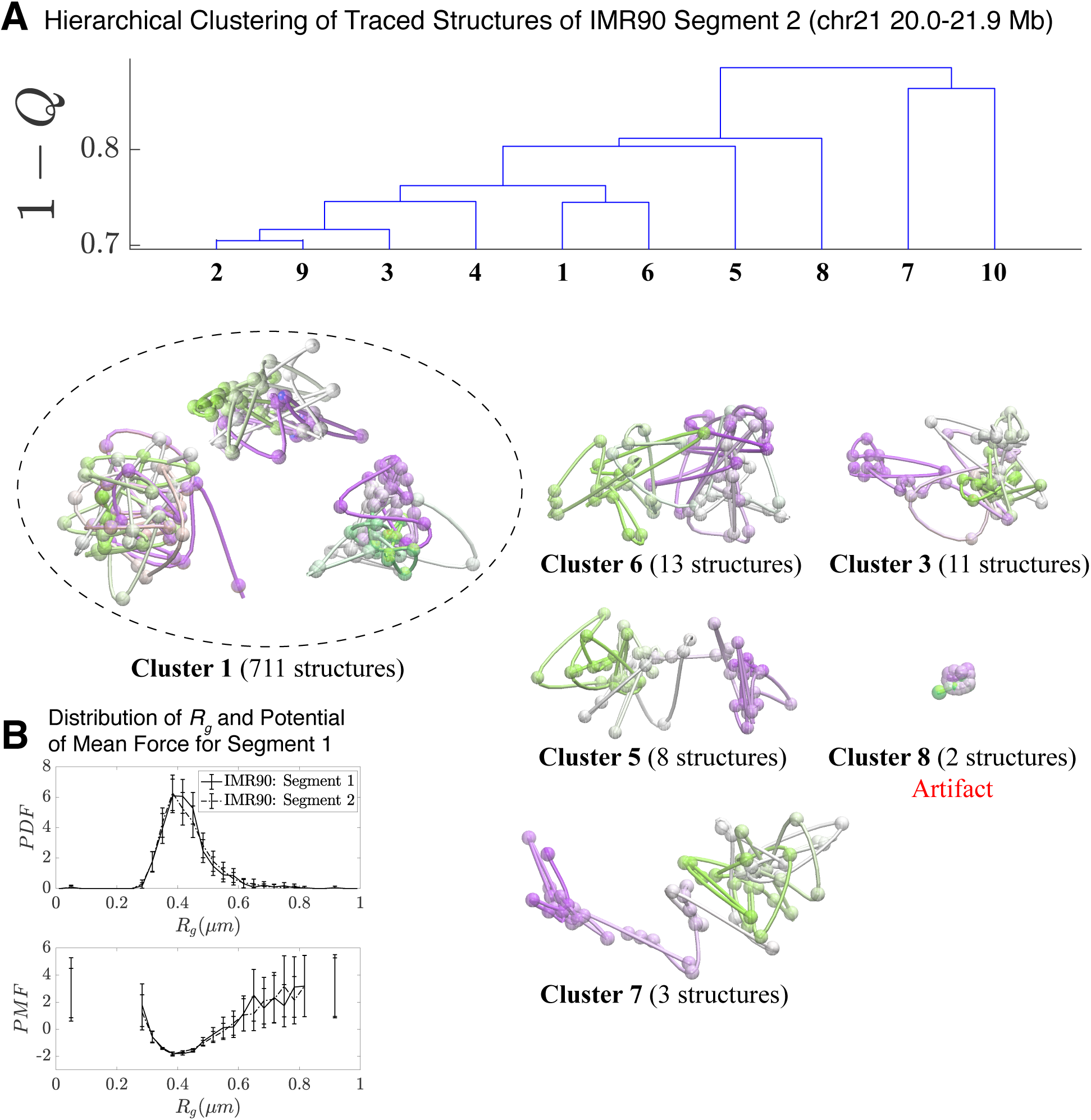
Hierarchical Clustering and the detailed structural analysis of traced Segment 2. (A) The dendrogram representation of the hierarchical clustering of Segment 2 (chr 21 20.0-21.9 Mb for IMR90 of [3]), where 1− *Q* is used as the distance between two structures. The dendrogram highlights 10 clusters—representative structures are shown for each of the featured clusters. Cluster 1 bears a close relation to the closed dumbbell structures observed for Segment 1 (Figure 2). Cluster 8 captures the highly dense chromatin structures that are attributed to experimental artifact; analogous to Cluster 3 in Figure 2. The additional clusters capture the gradual opening of Segment 2. However, a striking difference from the structures of Segment 1 occurs due to the lack of loop domains in Segment 2; as a result, the globular domains at the head and tail of the segment observed for Segment 1 do not exist. The lack of loop domains and domain boundaries leads to disordered structures that deviate from the open dumbbell structures observed for Segment 1. (B) The distribution of the Radius of Gyration (top) and the corresponding potential of mean force (bottom) are shown for the traced structures of Segment 1 and Segment 2 of IMR90. Both distributions exhibit a heavy tail to the right of the average value, indicating the existence of open, elongated structures.

**Figure S5.**
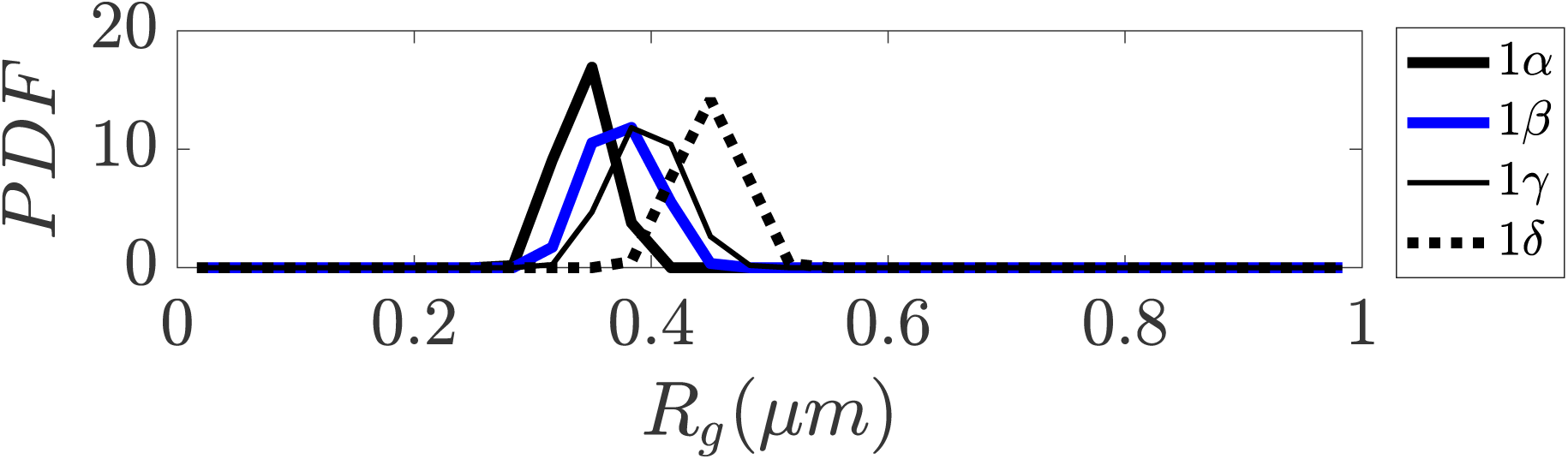
Distribution of Radius of Gyration for sub-clusters of closed dumbbell structures obtained from simulation. Sub-clusters of Cluster 1 in Figure 3 are denoted as 1α, 1β, 1γ, and 1δ. The sub-clusters of Cluster 1 characterize the gradual opening of the closed dumbbell structures observed in simulation.

**Figure S6.**
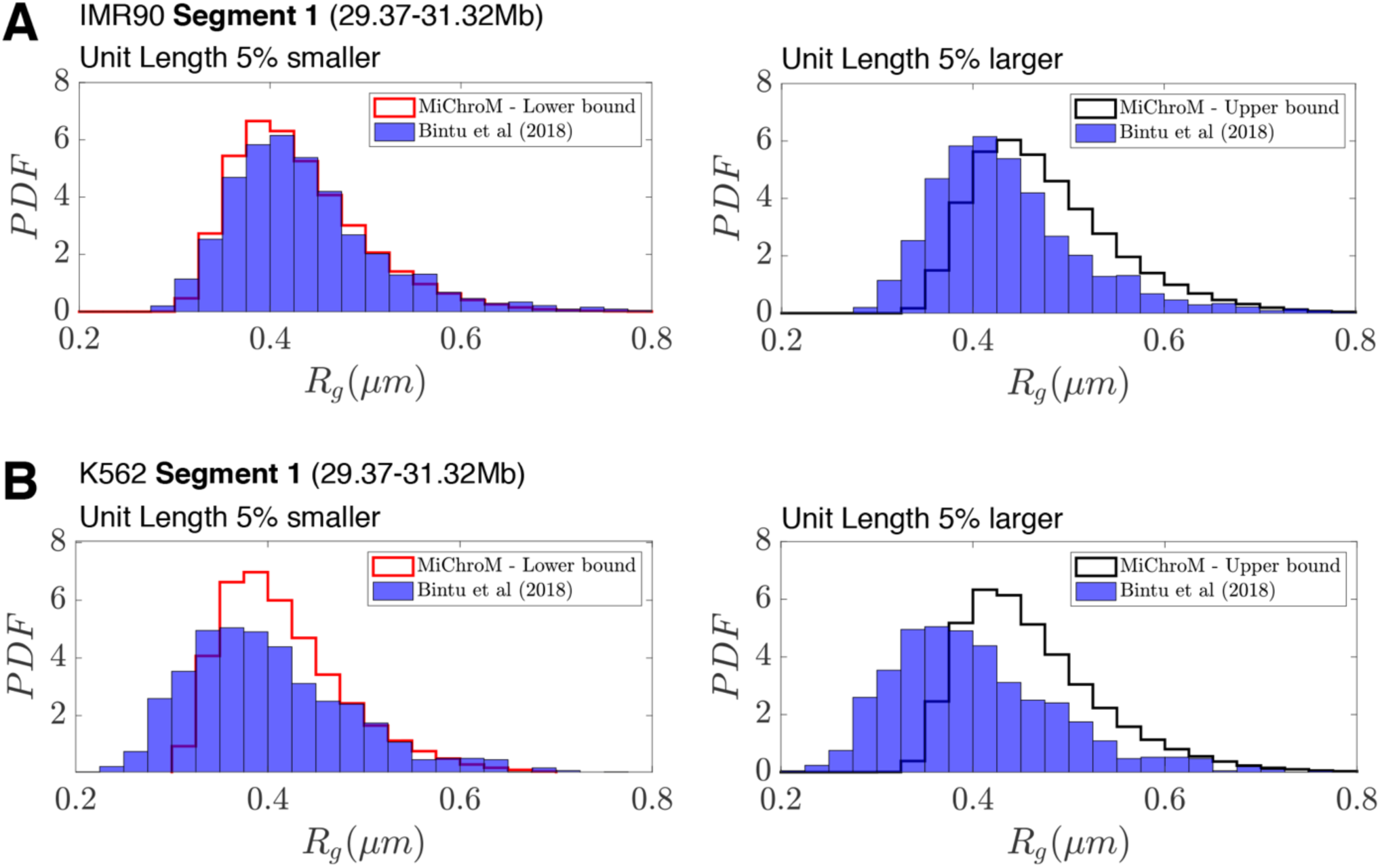
Deviations in the unit of length estimate can account for the differences in the experimental and simulated distributions of Radius of Gyration. (A) The distributions of Radius of Gyration for the experimentally traced IMR90 Segment 1 (A) and K562 Segment 1 (B) ([3]) are plotted alongside the distribution of Radius of Gyration from simulation when the unit length of our model (0.165μm) is 5% smaller or 5% larger.

**Figure S7.**
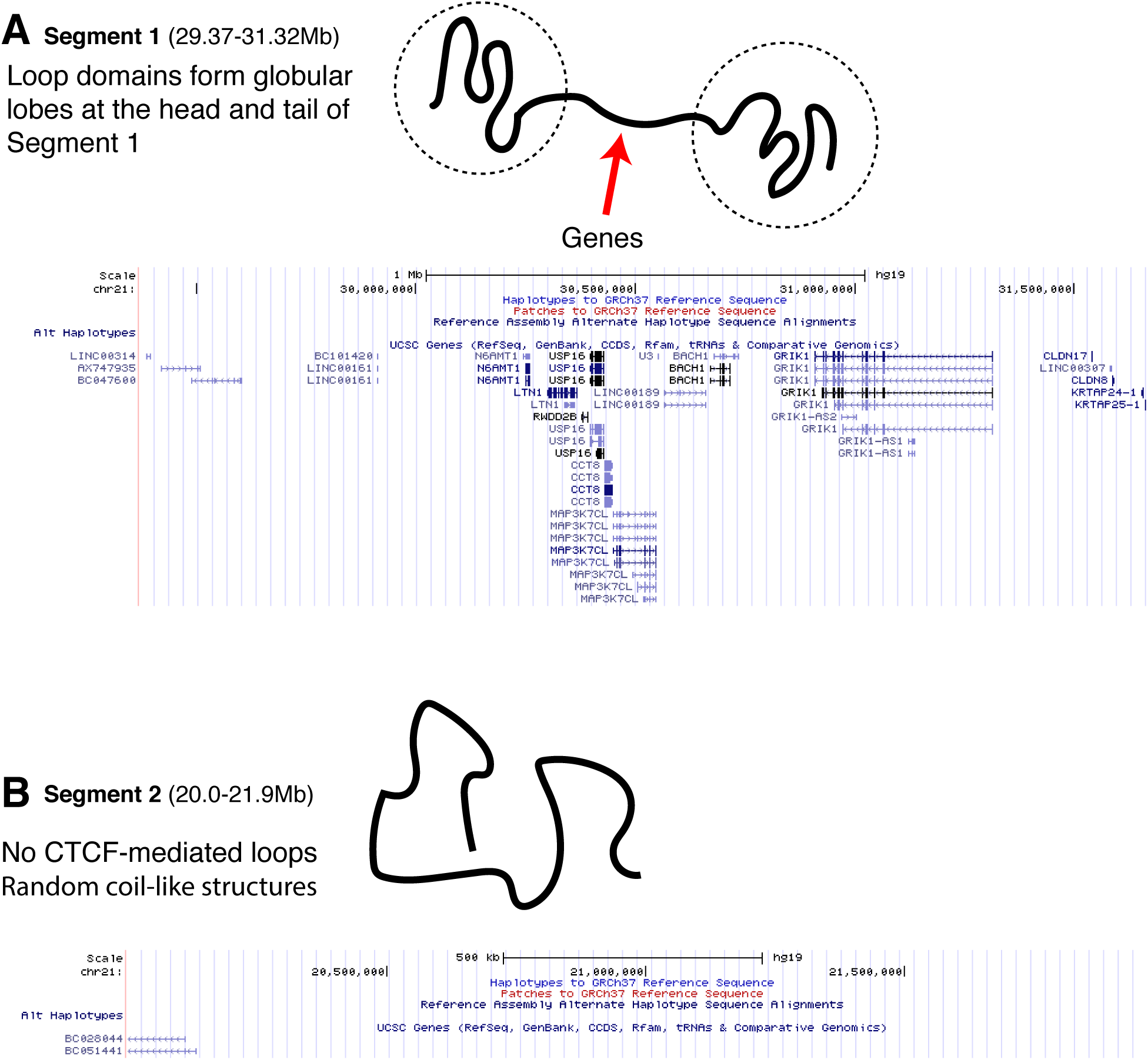
The positioning of genes along traced Segment 1 and Segment 2. The UCSC Genome Browser[4] is used to plot the UCSC Gene Track for (A) traced Segment 1 and (B) traced Segment 2. The positioning of the genes along Segment 1 appears primarily in the linker region sandwiched between the globular domains that are at the head and tail of the chromatin segment. Segment 2, which has no loop domains, also coincidentally has an apparent absence of genes.

## Methods

### MEGABASE

Maximum Entropy Genomic Annotations from Biomarkers Associated to Structural Ensembles (MEGABASE) was previously introduced in Ref: [5]. MEGABASE was trained to quantify the correlations between chromosome structural annotations (i.e., compartment annotations A1, A2, B1, B2, B3) with chromatin immunoprecipitation (ChIP-Seq) signals. This allowed for the inference of the chromatin types (compartment labels) for each 50 kb locus of chromatin, given information about the histone modifications present at that locus.

#### Discretization of ChIP-Seq Data Tracks

Chromatin Immunoprecipitatin (ChIP-seq) data was downloaded from ENCODE [6] for the cell lines explored in this manuscript: IMR90, HUVEC, K562, HMEC, H1-hESC, and HeLa-S3. We focused on 11 histone modification tracks: H2AFZ, H3K27ac, H3K27me3, H3K4me1, H3K4me2, H3K4me3, H3k79me2, H3K9ac, H3K9me3, and H4K20me1. These 11 tracks were previously shown to contain sufficient information to predict the chromosome structural ensembles for GM12878[5].

For each chromosome, the ChIP-Seq signal is re-casted into the data tracks at 50 kb resolution, i.e., loci of 50 kb in size. This is performed by summing the ChIP-Seq signal contained within each 50 kb locus respective of each experiment.

The integrated ChIP-seq signal for each 50 kb locus is assigned a discrete state ranging from 1 (low signal) to 20 (high signal). This is performed by creating a histogram for each experiment of the integrated signal for all of the 50 kb loci in the chromosomes of each cell type. All loci belonging to the top 5% of the distribution with the highest signal are assigned the highest signal state, i.e. 20. The remaining 19 signal states are defined by partitioning the remainder of the distribution linearly with respect to the signal strength; loci are assigned to those states according to their integrated signal.

#### Prediction of chromatin structural types from ChIP-Seq data using MEGABASE

The inferred probabilistic model ([5]) can be marginalized to predict the chromatin type for a given locus *l* when given the experimental ChIP-Seq measurements at loci *l*-2, *l*-1, *l, l*+1, and *l*+*2:*

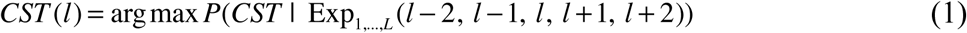

where *L*=11 is the number of epigenetic histone modifications used in this study. This allows for the prediction of the chromatin type (CST) for a given chromatin locus, given the ChIP-Seq signals for the 11 histone modifications at that locus. For additional details on the MEGABASE model, refer to Ref: [5].

### Minimal Chromatin Model (MiChroM)

The Minimal Chromatin Model (MiChroM) is a coarse-grained representation of individual chromosomes that was first introduced in Ref:[7]. The full energy function of MiChroM originally published in Ref:[7] is given by:

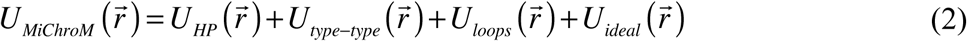

where

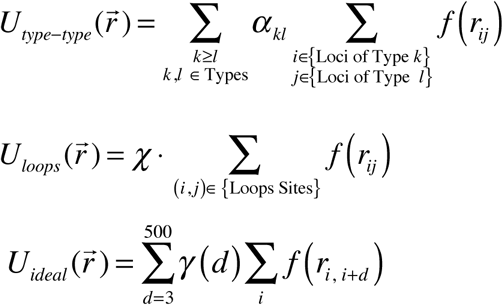

and

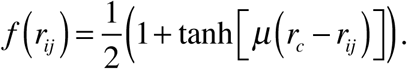

The first term *U*_*HP*_ is a homo-polymer potential that describes the connectivity (bonds and angles) between monomers—the monomers here represent a 50 kb span of DNA. The second term *U*_*type*−*type*_ describes the sequence-dependent interactions between pairs of monomers; this term captures the phase separation of chromatin loci into spatial compartments. The parameters ^*α*^ _*kl*_ describe the energetic stabilization when two loci of chromatin type *k* and *l* are spatially proximal. The third term *U*_*loops*_ describes the interaction between loop anchors that stabilize a CTCF-mediated loop. The final term *U*_*ideal*_ describes the local ordering in chromatin; a pair of loci are stabilized by an energy *γ* (*d*) that depends on the genomic distance between the loci pair, *d* = |*i* − *j*|.

The parameters *μ* = 3.22 and *r*_*c*_ = 1.78 were adjusted for the contact maps of GM12878 B-lymphoblastoid cells in dataset GSE63525 [1]. The parameters *α, χ*, and *γ* were iteratively trained [7] to be consistent with the DNA-DNA ligation map of chromosome 10 of human lymphoblastoid cells (GM12878)[1].

MiChroM considers 5 chromatin types A1, A2, B1, B2, B3 plus a non-specific type NA, which is used to label the centromere.

The α parameters, which govern the type-to-type interactions, are given in Table S1:

**Table S1.** MiChroM parameters for type-to-type interactions.

|  | A1 | A2 | B1 | B2 | B3 | NA |
| --- | --- | --- | --- | --- | --- | --- |
| A1 | -0.268028 | -0.274604 | -0.262513 | -0.258880 | -0.266760 | -0.225646 |
| A2 | -0.274604 | -0.299261 | -0.286952 | -0.281154 | -0.301320 | -0.245080 |
| B1 | -0.262513 | -0.286952 | -0.342020 | -0.321726 | -0.336630 | -0.209919 |
| B2 | -0.258880 | -0.281154 | -0.321726 | -0.330443 | -0.329350 | -0.282536 |
| B3 | -0.266760 | -0.301320 | -0.336630 | -0.329350 | -0.341230 | -0.349490 |
| NA | -0.225646 | -0.245080 | -0.209919 | -0.282536 | -0.349490 | -0.255994 |

The parameter *χ* governing the loop interactions is equal to −1.612990.

The ideal chromosome potential is given by:

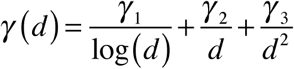

with parameters *γ* _1_ = −0.030, *γ* _2_ = −0.351, *γ* _3_ = −3.727.

The reduced MiChroM energy function used in this manuscript omits CTCF-mediated loops unless stated otherwise:

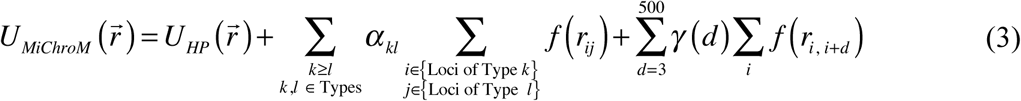

For comparison with the DNA-tracing structures of Bintu et al[3], simulations of chromosome 21 for cell types IMR90 and K562 with CTCF-mediated loops were generated using the full energy function of MiChroM.

## Notes

### Competing Interest Statement

The authors have declared no competing interest.

### Summary of Updates

Figure 2E was added to show the positioning of the genes. The experimental 3D structures examined in this manuscript (of Bintu et al, Science, 2018) were erroneously referred to as obtained via "super-resolution microscopy". The manuscript has been corrected to reflect that those structures were obtained using diffraction-limited methods. Several additional typos were fixed, including "Genonic" -> "Genomic" in Figures 1 and S1. The introduction text was reorganized. The Supporting Information was appended at the end of the manuscript file.

